# Locus-Scale Massively Parallel Reporter Assays

**DOI:** 10.64898/2026.07.24.740649

**Authors:** Abby V. McGee, Carina G. Biar, Beth K. Martin, Tony Li, Jean-Benoît Lalanne, Haedong Kim, Jay Shendure

## Abstract

Interactions among *<u>c</u>is*-regulatory <u>e</u>lements (CREs) are central to mammalian gene regulation. <u>L</u>ong <u>@</u>$$ <u>m</u>assively <u>p</u>arallel reporter <u>a</u>ssays (LAMPRAs) integrate combinatorial cloning, molecular barcoding and long- and short-read sequencing, to scalably measure how CRE identities, numbers, spacings, orders, orientations, and interactions shape regulatory output at multi-kilobase length scales. As a proof-of-concept, we assay 36,000 *×* 5-kb <u>s</u>ynthetic *<u>c</u>is-*regulatory loci (sCRLs), each a 5 *×* 1-kb random combination of enhancers, insulators and spacers, to model how locus composition drives gene expression.

## MAIN TEXT

MPRAs have transformed our ability to study gene regulation by enabling the multiplex functional analysis of thousands of candidate CREs per experiment^1–3^. However, the lengths of CREs assayed in MPRAs have overwhelmingly been shaped by the technical limitations of microarray-based DNA synthesis rather than biological considerations, such that nearly all MPRA studies to date assay sub-300 bp fragments^4^ positioned closely upstream^1–3^ or downstream^5^ of a minimal promoter.

Although straightforward to implement, conventional MPRAs are poorly matched to the architecture of mammalian gene regulation in at least three ways: (i) <u>length</u>: *in vivo-*validated developmental enhancers are overwhelmingly >300 bp^6,7^, while “super-enhancers” can span kilobases^8,9^; (ii) <u>distance</u>: enhancer-promoter distances range from ∼10^1^ to ∼10^6^ base-pairs^10–12^; and (iii) <u>combinatorial logic</u>: regulatory output is shaped by interactions among multiple kinds of CREs (*e.g.* enhancers, promoters, insulators, silencers, facilitators, etc.) in a manner that depends on their identities, numbers, spacings, orders, orientations, and interactions^13–17^. These mismatches highlight a locus-scale regime of multi-element, kilobase-to-megabase-scale regulatory grammar that conventional MPRAs are poorly equipped to probe.

Although deep learning is proving highly effective for modeling the regulatory grammar of cell type-specific enhancers by training on genomewide biochemical profiles^18–20^, we are skeptical that analogous efforts to model locus-scale regulatory grammar^19,21^ will reach their potential without fundamentally new kinds of data. Enhancer grammar is tractable because: (i) chromatin accessibility is a reasonable, albeit imperfect, proxy for enhancer activity^22,23^; and (ii) evolution has generated thousands of independent training examples per cell type^24^. Neither advantage holds at the locus scale. Although methods like Hi-C quantify 3D contact frequencies of enhancer-promoter pairs in support of useful heuristics (*e.g.* “activity-by-contact”)^25,26^, 3D proximity tracks poorly with functional enhancer-promoter communication^27^. Furthermore, because evolution varies enhancer sequence much more freely than locus architecture, the arrangements that would let a model fully disentangle locus-scale grammar (*e.g.* the identities, numbers, spacings, orders, orientations, and interactions of CREs, each varying independently) are largely absent from any mammalian genome, however many we sequence.

Consequently, major gaps remain in our understanding of *cis*-regulatory logic beyond the level of fragmented candidate enhancers. There have been some successful efforts to extend MPRAs to longer sequences (*e.g.* libraries derived from shearing or digesting genomic DNA^5,28^; multiplex assembly of short oligonucleotides^29^) but these remain technically demanding and comparatively low-throughput. Combinatorial MPRAs probing defined pairs or triplets of CREs offer more controlled interrogation of interactions^29–35^. NeMECiS, for example, exhaustively fuses triplets of ∼200 bp fragments into ∼600 bp synthetic CREs, enabling the authors to shed light on how intra-CRE regulatory modules combine to shape CRE activity in response to a graded developmental signal^35^. To date, however, such efforts have remained at sub-kilobase length scales, on par with individual enhancers rather than genomic loci. Studies leveraging genomic integration into landing pads with FACS-based readouts extend the MPRA concept to greater enhancer-promoter distances, but at the cost of throughput and generality, as the landing pads are positioned within only one or a few endogenous genomic contexts^36^. Collectively, these methods and studies underscore the importance of locus-scale regulatory grammar, as well as the dearth of methods to interrogate it in a systematic, scalable manner.

LAMPRAs aim to fill this gap, by using modular, slot-based combinatorial assembly^37–39^ to generate multi-kilobase <u>s</u>ynthetic *<u>c</u>is-*regulatory loci (sCRLs) (**Fig. 1a**). Briefly, a CRE library is synthesized with identical flanking Type IIS restriction sites and bidirectionally cloned to a set of slot-specific intermediate vectors (**Supplementary Fig. 1a**). Slot libraries are then combinatorially assembled and cloned upstream of a fixed reporter cassette composed of a minimal promoter (minP), degenerate barcode, and GFP-encoding open reading frame (minP-BC-GFP). Because of the 1:1 correspondence between intermediate vectors and slots, distinct CRE libraries can (optionally) be directed to specific slots. The barcode uniquely tags each construct, enabling barcode-locus association by long-read sequencing^40,41^. The barcoded sCRL library is randomly integrated to the genome via piggyBac transposition, providing chromatinized context while also retaining the potential to accommodate larger sCRL lengths than lentiviral or safe harbor delivery^42^. Finally, sCRL activity is quantified by measuring DNA-normalized RNA barcode abundance by short-read amplicon-sequencing (**Fig. 1b**).

**Figure 1.**
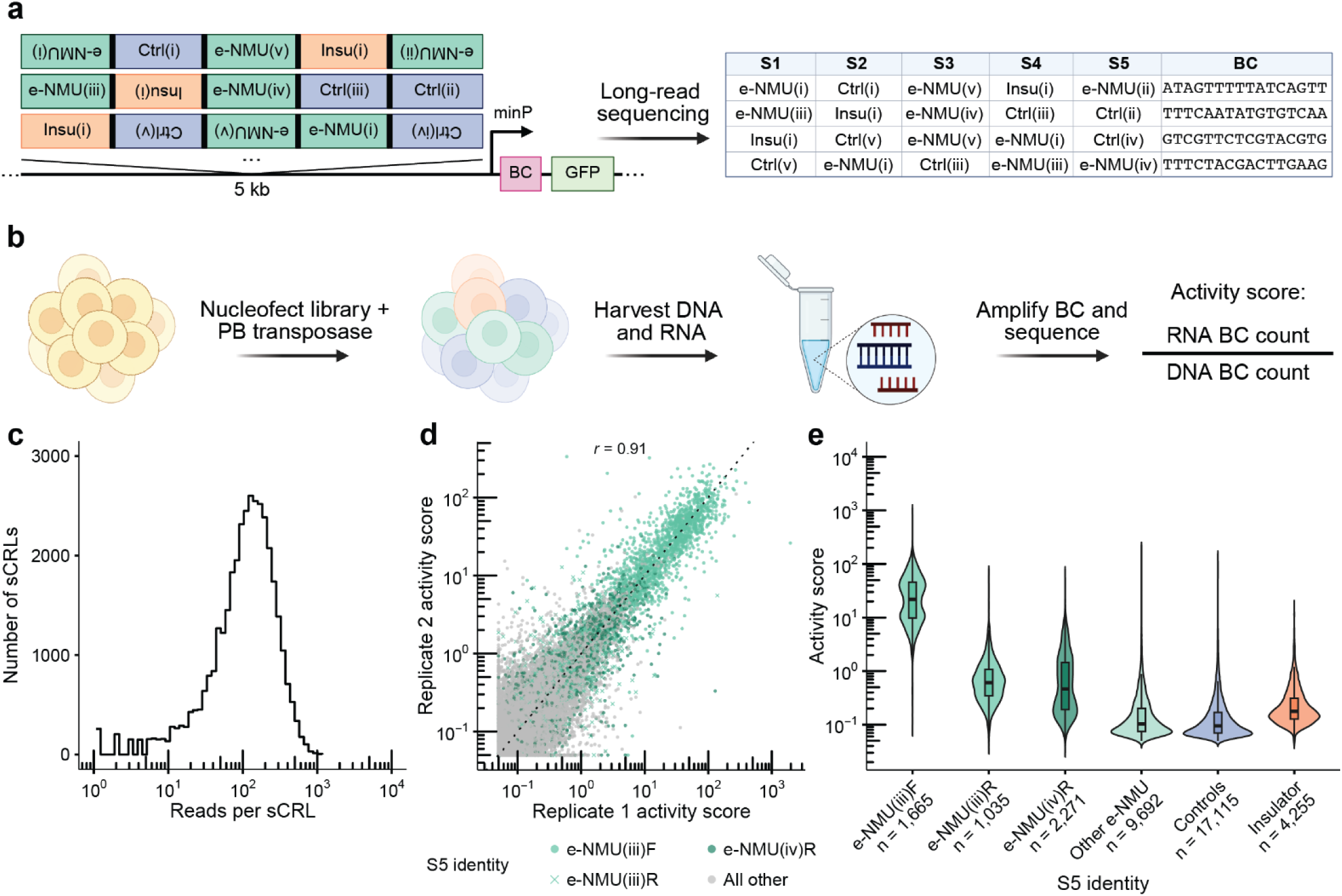
<u>L</u>ong <u>@</u>$$ <u>m</u>assively <u>p</u>arallel reporter <u>a</u>ssays (LAMPRAs) enable the functional characterization of kilobase-scale synthetic *cis*-regulatory loci (sCRLs). **(a)** Schematic of LAMPRA, as applied to 11 *×* 1-kb sequences, including 5 NMU candidate enhancers (e-NMU), 5 dinucleotide-matched spacers (Ctrl), and 1 synthetic insulator (Insu). Combinatorial, bidirectional cloning of these elements to a 5-slot LAMPRA vector was followed by introduction of a 16-bp degenerate barcode (BC) into the 5’ UTR of a minimal promoter (minP)-GFP cassette. Long-read sequencing (PacBio) was used to associate each of the resulting 5 *×* 1-kb sCRLs with specific barcode(s). **(b)** The e-NMU LAMPRA library was randomly integrated into the genome of K562 cells, in triplicate, via nucleofection and piggyBac transposition. This was followed by the conventional MPRA workflow, including harvesting of nucleic acids, amplicon sequencing of barcodes derived from both RNA and DNA, and calculation of activity scores. **(c)** Distribution of sequencing coverage per sCRL upon PacBio sequencing of the cloned e-NMU LAMPRA library. Histogram shows the number of sCRLs (y-axis) covered at various depths (x-axis, log10 scale; tick labels indicate untransformed read counts). Bins are equally spaced on a log10 scale, and all sCRLs supported by at least one mapped read are included. The distribution is approximately log-normal, with a left tail of low-coverage sCRLs. **(d)** Correlation of log-transformed activity scores between a representative pair of biological replicates (independent nucleofections and piggyBac integrations of the same library in K562 cells). Each point (n = 35,567) is a unique sCRL detected in ≥2 DNA UMIs in both plotted replicates, stratified by S5 identity. “All other”, sCRLs with any element besides e-NMU(iii)F, e-NMU(iii)R, or e-NMU(iv)R in S5. For these, minP is the effective promoter. Dashed line, y = x. **(e)** Violin plots of the distribution of activity scores (y-axis, log10 scale) for sCRLs stratified by S5 identity, sorted by mean activity. Overlaid boxes denote the median and interquartile range; whiskers extend to 1.5× IQR. “Other e-NMU”, sCRLs with any e-NMU element other than e-NMU(iii)F, e-NMU(iii)R, or e-NMU(iv)R in S5; “Controls”, all sCRLs with Ctrl(i)-(v) in S5; “Insulator”, all sCRLs with Insu(i) in S5.

For this proof-of-concept, we focused on five distal candidate enhancers of the human *NMU* gene that reside 30-98 kb upstream of its transcriptional start site (TSS), which we initially identified by CRISPRi screening in K562 cells and named e-NMU (i) through (v)^11^. These broadly correspond to F1, F2, e1, e2, and F3 in the nomenclature of Zhou *et al.*^17^, but are centered on coordinates defined by Gasperini *et al.*^11^. Altogether, we designed eleven 1-kb elements, corresponding to the e-NMU regions (n = 5), spacers obtained by dinucleotide-matched shuffling of the e-NMU regions (n = 5), and a synthetic insulator that concatenates five documented insulators^43^ (n = 1) (**Supplementary Table 1**).

We obtained these eleven 1-kb elements via commercial synthesis (IDT) and subjected them to a “five-slot” LAMPRA. Although our scheme allows for distinct libraries to be cloned to each slot (**Supplementary Fig. 1a**), here a single library (11 elements × 2 orientations) was cloned to all five intermediate vectors. Following combinatorial assembly, introduction of a degenerate barcode, barcode-construct association (PacBio Vega), and QC filtering, we obtained 36,922 uniquely barcoded 5-kb sCRLs with sufficient representation for downstream analyses, out of 5.2 million possibilities ([11 elements × 2 orientations] ^ 5 slots). Given a ∼$1,000 synthesis cost per 1-kb element, this equates to $0.30 per sCRL assayed -- an upper bound, as synthesis is a one-time cost and deeper sampling of the 5.2-million sCRL space adds only cheap assembly and transformation. The eleven elements were reasonably well-balanced with respect to abundance (7.3-fold range) and orientation (51% forward, 49% reverse) (**Supplementary Fig. 1b**), and the 5-kb sCRLs exhibited reasonable uniformity as well (59% within 4-fold, 79% within 10-fold) (**Fig. 1c**; **Supplementary Fig. 1c**). On average, each sCRL contained 1.95 enhancers, 2.45 spacers, and 0.60 insulators, and was represented by 3.14 independent barcodes. Correspondingly, 8%, 3%, and 51% of sCRLs lacked any enhancer, spacer, or insulator, respectively (the latter due to our inclusion of only a single insulator).

We adopt the following nomenclature: the five NMU enhancer regions are named e-NMU(i) to e-NMU(v)^11^, their matched controls are named Ctrl(i) to Ctrl(v), and the synthetic insulator is named Insu(i) (**Supplementary Table 1**). Element orientation is indicated by an F or R following the element name, while slots are designated S1-S5 in the orientation of the minP-BC-GFP cassette (**Supplementary Fig. 1a**). For example, e-NMU(iv)R-S1 references the reverse orientation of the e-NMU(iv) candidate CRE in the slot furthest from the minP-BC-GFP cassette.

We integrated the e-NMU LAMPRA library into K562 cells in triplicate by piggyBac transposition, grew the cells for 10 days to dilute out unintegrated episomes, and then harvested genomic DNA and total RNA. Following amplification and sequencing, MPRA activity scores were calculated as the ratio of RNA to DNA barcode abundance. Activity scores were obtained for 36,033 of 36,922 (98%) sCRLs in ≥2 replicates (min. 2 DNA UMIs to calculate an activity score; mean 549 +/- 490 DNA UMIs per sCRL per replicate). As transfection replicates were well correlated (Pearson *r* = 0.91 for pairwise comparisons of log-transformed activity scores; **Fig. 1d**), all downstream analyses are based on the mean activity score for each sCRL across replicates.

The activity scores of these 36,033 sCRLs spanned nearly three orders of magnitude (range 0-669; **Fig. 1d-e**). However, on cursory inspection, much of this variation was attributable to sCRLs harboring e-NMU(iii)F-S5 (*i.e.* the forward orientation of the e-NMU(iii) candidate enhancer in the promoter-proximal slot). The mean activity score of these sCRLs was 32.7 (s.d. 38.2; range 0.07-669; n = 1,665), versus 0.296 (s.d. 1.62; range 0-85; n = 34,368) for all remaining sCRLs, a 110-fold difference (**Fig. 1e**). On average, sCRLs bearing e-NMU(iii)R-S5 or e-NMU(iv)R-S5 also exhibited elevated activity, albeit less so than the e-NMU(iii)F-S5 subset (**Fig. 1e**). All other S5 identities, including e-NMU(iv)F-S5, the other e-NMU elements, controls and the insulator, were unremarkable, exhibiting similarly low-mean, long-tailed distributions (**Supplementary Fig. 1d**).

Because e-NMU(iii)F, e-NMU(iii)R, and e-NMU(iv)R exhibit substantially elevated, orientation-dependent activity when positioned in S5, we reasoned that in these subsets of sCRLs, they serve as the effective promoter, driving transcription of the barcoded reporter strongly enough to overwhelm the comparatively weak minP. This motivated us to recast the full dataset in terms of four promoter groups, each defined by its effective promoter: the three subsets in which one of these elements occupies S5 (e-NMU(iii)F-S5: n = 1,665; e-NMU(iii)R-S5: n = 1,035; e-NMU(iv)R-S5: n = 2,271), and a fourth, "minP" group comprising all remaining sCRLs, in which minP remains the dominant promoter (n = 31,062).

A consequence of this framing is that the number of enhancer-occupiable slots differs by group. In the three alternative-promoter groups the promoter occupies S5, leaving S1-S4 (up to four positions) for upstream e-NMU enhancers, whereas in the minP group the promoter resides in the fixed downstream cassette and all five slots (S1-S5) are available. We therefore assess enhancer dose-response over the relevant range for each group (**Fig. 2a**). Baseline activity in the absence of any upstream e-NMU element varied ∼130-fold across the four promoter groups (e-NMU(iii)F-S5: 7.85; e-NMU(iii)R-S5: 0.62; e-NMU(iv)R-S5: 0.15; minP: 0.061), and mean activity rose with the copy number of upstream e-NMU elements for all four groups (**Fig. 2a**; **Supplementary Fig. 2a**).

**Figure 2.**
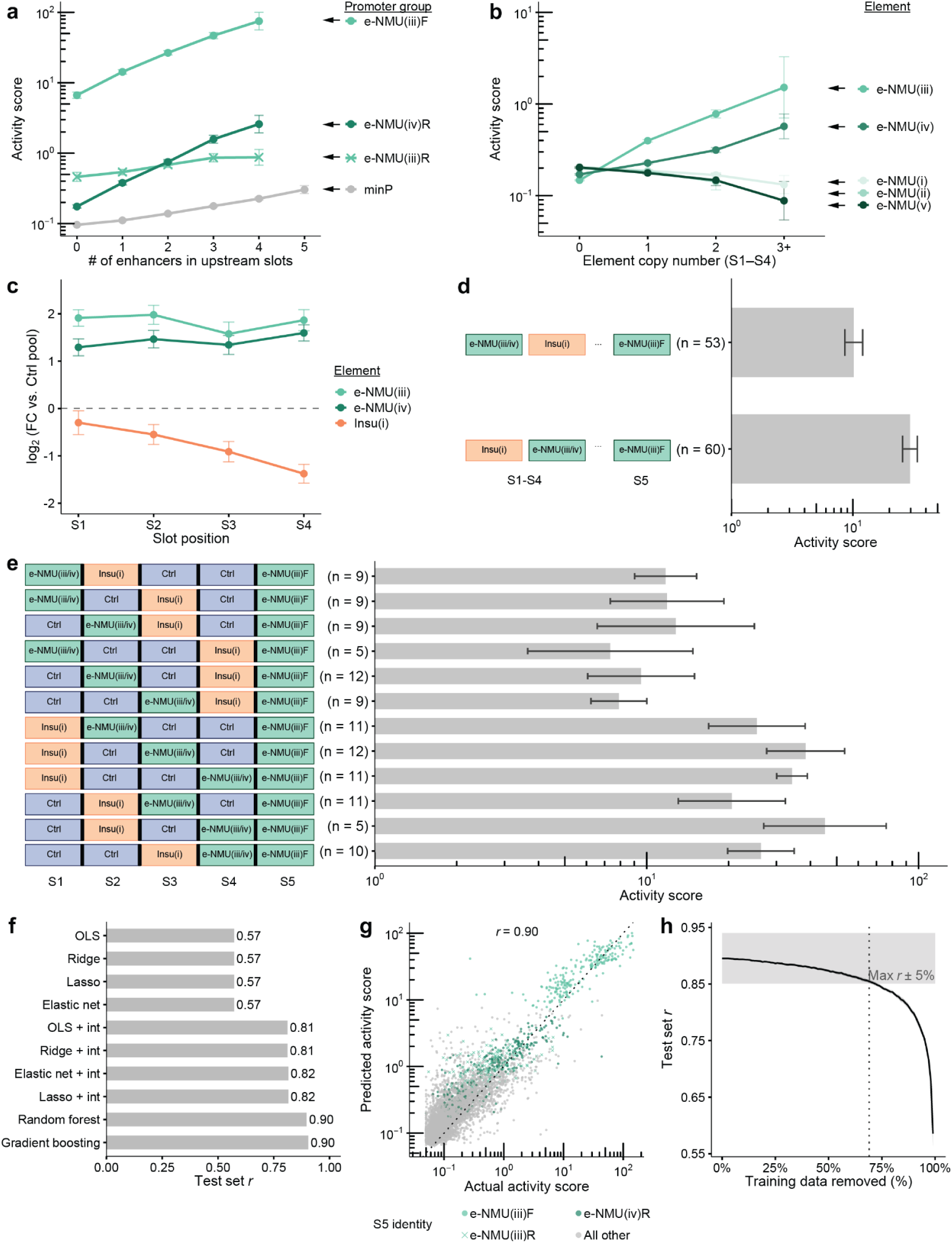
A multiplicative, promoter-dependent model relates sCRL composition to activity, with enhancers and insulators acting through distinct positional logic. **(a)** Mean LAMPRA activity score, plotted on log-scale, as a function of the number of e-NMU enhancer elements upstream of the effective promoter (n = 36,033 sCRLs). Elements are stratified to each of four promoter groups. Error bars, 95% confidence intervals (t-distribution). **(b)** Mean LAMPRA activity score, plotted on log-scale, as a function of the number of instances of a given e-NMU element across slots S1-S4. Error bars, 95% confidence intervals (t-distribution). **(c)** Log_2_ fold-difference in mean activity between pooled sCRLs bearing e-NMU(iii), e-NMU(iv), or Insu(i) at each slot (S1-S4) vs. pooled sCRLs bearing Ctrl(i)-(v) at the same slot. This analysis is restricted to the e-NMU(iii)F-S5 promoter group (e-NMU(iii): n = 294; e-NMU(iv): n = 524; Insu(i): n = 620), and enhancer trends are computed from insulator-free sCRLs. Dashed line indicates no fold-difference. Error bars, 95% confidence intervals (±1.96 x SE). **(d)** Horizontal bar plot of mean activity score (x-axis, log10 scale) for sCRLs with a single active e-NMU element (e-NMU(iii) or (iv), in either orientation) positioned either upstream or downstream of a single Insu(i) element within S1-S4 (e-NMU(iii)F-S5 subset; sCRLs with any inactive e-NMU element in S1-S4 excluded, n = 113). Error bars, 95% confidence intervals (t-distribution). **(e)** Similar to panel **d**, but here broken down by full sCRL architecture. Diagrams on left indicate the position of the Insu(i) and the active e-NMU element within slots S1-S4. Error bars, 95% confidence intervals (t-distribution). **(f)** Bar chart comparing Pearson’s *r* on held-out activity scores across all evaluated models, sorted by performance on the test set (n = 5,405); *r* values are indicated at the end of each bar. OLS: ordinary least squares. “+ int” indicates inclusion of interaction terms. **(g)** Scatter plot of predicted vs. actual LAMPRA activity scores (both axes on log10 scale) for the most generalizable model by holdout Pearson’s *r* across S5 identities (random forest), stratified by S5 identity (n = 5,405). “All other”, sCRLs with any element besides e-NMU(iii)F, e-NMU(iii)R, or e-NMU(iv)R in S5. For these, minP is treated as the effective promoter. Dashed line, y = x. **(h)** Random forest performance (Pearson’s *r*) by training data fractions. Gray box spans ±5% of the *r* obtained with the complete training set (n = 27,565 sCRLs); dashed line marks the smallest training set whose lower 95% confidence bound recovers 95% of the full-training-set *r*. Solid line, mean across 50 downsampling permutations. Gray ribbon, 95% confidence intervals (t-distribution).

Having established that upstream e-NMU enhancer dose scales activity within each promoter group, we sought to determine which of the five e-NMU elements drive this response. Increasing the copy number of e-NMU(iii) or e-NMU(iv) in S1-S4 raised LAMPRA activity, whereas e-NMU(i), (ii), and (v) showed no such dose-response (**Fig. 2b**). We therefore designate e-NMU(iii) and e-NMU(iv) as active enhancers for all downstream analyses. However, these are the same two elements that act as orientation-specific promoters when positioned in S5, raising the question of whether their activity in S1-S4 reflects genuine enhancer function or residual promoter activity.

Two observations argue for the former. First, activity increased log-additively with the copy number of the two active enhancer elements but not the two inactive ones (**Supplementary Fig. 2b**). Second, e-NMU(iii) and e-NMU(iv)’s contributions to sCRL activity are orientation-independent in S1-S3, but orientation-dependent in S5 (**Supplementary Fig. 2c-d**). Because promoter activity is directional whereas enhancer activity is classically independent of orientation, the loss of orientation-dependence upon relocation from S5 to upstream slots supports a bona fide enhancer function distinct from these elements’ promoter activity in S5.

Overall, these data are consistent with a multiplicative model^32,33,35^ in which, within each promoter group, log-activity is set by two terms: (i) a promoter-specific baseline (**Fig. 2a**); and (ii) a dose-dependent contribution from active, upstream enhancers (**Fig. 2b**). The per-copy enhancer gain (*i.e.* the slope of this log-additive dose-response) is also promoter-specific (e-NMU(iii)F-S5: 1.87; e-NMU(iii)R-S5: 1.22; e-NMU(iv)R-S5: 2.04; minP: 1.25; **Fig. 2a**; **Supplementary Fig. 2a**), and, notably, does not track with the promoter-specific baseline. For example, e-NMU(iii)R-S5 pairs a comparatively high baseline with one of the shallowest gains, whereas e-NMU(iv)R-S5 pairs a much lower baseline with one of the steepest gains. Baseline activity and enhancer-responsiveness thus appear to be promoter-specific and independent (*i.e.* knowing one does not predict the other).

To quantify the impact of the insulator element, Insu(i), we focused on the highest-activity promoter group (e-NMU(iii)F-S5 sCRLs; n = 1,665), reasoning that its high dynamic range affords the most power to detect positional effects. Within this subset, the two active enhancers behaved in a distant-invariant manner within the 1-5 kb range, increasing activity by ∼1.83-fold (e-NMU(iii)) and ∼1.43-fold (e-NMU(iv)) (**Fig. 2c**). In contrast, sCRLs bearing one or more Insu(i) elements were >2-fold less active than those bearing none (**Supplementary Fig. 3a**), and this effect was strongly position-dependent, with larger decreases for insulators closer to the promoter (**Fig. 2c**). We hypothesized that rather than insulators themselves having an intrinsic positional dependency, promoter-proximal insulators are simply statistically more likely to interpose between an enhancer and the promoter than promoter-distal ones. To test this, we examined e-NMU(iii)F-S5 sCRLs containing exactly one insulator and one active e-NMU enhancer in S1-S4 (n = 113), and found 2.9-fold higher activity when the insulator lay upstream of the enhancer than when it intervened between enhancer and promoter (**Fig. 2d-e**; **Supplementary Fig. 3b-c**). This effect was independent of the insulator’s orientation (**Supplementary Fig. 3d**). Thus, within a single locus, e-NMU enhancers and the synthetic insulator act through fundamentally different positional logics: enhancers contribute similarly regardless of where they sit upstream, whereas the insulator suppresses activity specifically when interposed between an enhancer and its target promoter. Although these behaviors align with textbook definitions of enhancers^44^ and insulators^45,46^, they have rarely been revisited in a systematic, controlled fashion, and their clean recovery here validates the LAMPRA framework.

Altogether, these composable properties suggest that relatively simple techniques should be effective for modeling these data. To test this, we trained a series of models to predict activity scores as a function of sCRL composition (**Fig. 2f**), in which each sCRL was featurized by the identity and orientation of the element at each slot. All models leveraged sCRLs present in ≥2 replicates (n = 36,033), with 15% set aside for model evaluation (n = 5,405), and the remainder split into training (90%; n = 27,565) and validation (10%; n = 3,063) sets. Linear models with main effects only achieved *r* = 0.57, regardless of regularization (OLS, ridge, lasso, elastic net). Adding pairwise interaction terms raised performance substantially (*r* = 0.81-0.82), implicating element interactions as a meaningful contributor to sCRL activity. Tree ensembles performed best (random forest and gradient boosting, *r* = 0.90), suggesting additional structure beyond pairwise interactions. Importantly, this high overall *r* is partially attributable to trivially predicted differences between promoter groups. Applying random forest modeling to each promoter group separately, we achieved the best performance on the e-NMU(iii)F-S5 sCRLs, which have the largest dynamic range (*r* = 0.81), followed by the large minP group (*r* = 0.56) then e-NMU(iv)R-S5 (*r* = 0.47) and e-NMU(iii)R-S5 (*r* = 0.39) (**Fig. 2g**). Of note, a downsampling analysis showed that 31% of the training data was sufficient to recover 95% of the maximum Pearson *r* (**Fig. 2h**), suggesting that more complex libraries will be not only measurable but also modelable with this framework.

In summary, LAMPRA extends the classic MPRA framework to the locus scale, combining combinatorial assembly, barcoding, and long- and short-read sequencing. Applied to ∼36,000 × 5-kb sCRLs of candidate e-NMU elements, we resolved regulatory output into three separable quantities: a promoter-specific baseline, a promoter-specific responsiveness to enhancer dose (the per-copy gain of a log-additive dose-response), and an enhancer-intrinsic effect set by element identity rather than orientation or position. Notably, promoter baseline and responsiveness varied independently, at least within the small set of promoters sampled here. Our findings accord with a recent preprint from Tan *et al.*, who, assaying ∼25,000 enhancer–promoter pairs, similarly find responsiveness to be an intrinsic promoter property independent of basal output, with output scaling with enhancer activity in a multiplicative, power-law manner whose exponent is promoter-specific^47^. That two different assays converge on baseline-independent responsiveness argues it is a general feature of enhancer-promoter interactions, albeit both in K562, a single cancer cell line.

The active elements e-NMU(iii) and e-NMU(iv) correspond to the “e1” enhancer and “e2” facilitator identified by Lis and colleagues^17,48^. The e1 enhancer harbors divergent TSSs at a putative LTR boundary, consistent with e-NMU(iii) acting in S5 as a strong promoter in one orientation (high baseline, high gain) and a weaker one in the other (moderate baseline, low gain). Of note, although Lis and colleagues identified no TSS in the orientation of e2^17,48^ matched to e-NMU(iv)R-S5, this orientation and positioning of e-NMU(iv) nonetheless defined its own promoter group, the lowest baseline of the four, yet the steepest enhancer responsiveness, potentially reflecting the copy-number-dependent activity observed by Lis and colleagues when two otherwise inactive e2 elements were fused in tandem^17^. We suspect e-NMU(iv)’s contribution reflects direct promoter activity in some architectures and position-dependent enhancer/facilitator activity in others, a distinction that will require further experiments to resolve.

Whether minP is a bona fide promoter has been questioned, along with the broader concern that signal in conventional MPRAs may partly reflect eRNA production or cryptic initiation within CREs rather than canonical promoter activity^28,47,49^. Our data do not resolve this, but the minP group, whatever the molecular origin of its signal, is the lowest-baseline group, effectively overridden for all sCRLs where a promoter-capable element resides in S5. The slot-addressability built into LAMPRA (available but not exploited here) offers a direct remedy, as eschewing minP and instead restricting S5 to a a defined library of endogenous or designed promoters would raise sensitivity and turn promoter identity into a systematically varied dimension, enabling direct dissection of the determinants of responsiveness to elements and combinations of elements within 5 kb. Upstream slots (S1-S4) would remain free for enhancers, insulators, and other kinds of CREs, supporting rich exploration of additivity, interaction, and positional logic.

Particularly at longer length scales, deep learning models of gene regulation are now limited by data rather than GPUs. LAMPRA’s combinatorial, locus-scale architecture is well suited to generating the training sets such models require. Moreover, the modular slot architecture generalizes well beyond this proof-of-concept, as distinct element libraries (*e.g.* enhancers, promoters, synthetic circuit parts, or gene bodies) can be addressed to specific slots^38,50^; constructs >5 kb are reachable through sequential combinatorial cloning and deliverable via piggyBac; and libraries scale readily in slot number, element length and combinatorial complexity. Beyond generating training data, these systematic, high-throughput measurements could also enable more data-driven definitions and characterizations of emerging element classes such as facilitators and super-enhancers⁴⁸⁻⁵⁰.

Underlying this opportunity is a deeper point, in that endogenous sequence alone may be insufficient to resolve regulatory grammar at longer length scales, because natural loci reflect evolutionary history rather than an independent sampling of the possible. The activity-by-contact heuristic^26^ is illustrative, as where Hi-C is unavailable, genomic distance substitutes for contact and works surprisingly well, though it is unclear whether enhancer activity genuinely decays with distance or whether greater distance simply raises the chance that a boundary element intervenes. Our own data favor the latter, as the synthetic insulator suppressed activity when interposed between enhancer and promoter, rather than as a function of distance alone. More broadly, as evolution has diversified enhancer sequences more rapidly than locus architectures, even sequencing and functionally characterizing every extant mammalian genome in every cell type may prove insufficient. In contrast, LAMPRA can construct and test vast numbers of locus architectures directly.

## ACKNOWLEDGEMENTS

We are grateful to members of the Shendure Lab, particularly Kshitij Rai, Sophie Seidel, Aidan Keith, Sanjay Kottapalli, Jonas Koeppel, Olga Oseth, Lily Bastian and the gene regulation subgroup, as well as to colleagues Nadav Ahituv and Martin Kircher and members of their labs, for technical advice, comments, suggestions, and discussions on this work. We acknowledge the use of Biorender.com to generate schematics for many figures in this manuscript. This work was supported by the Seattle Hub for Synthetic Biology, a collaboration between the Allen Institute, Biohub and University of Washington (CZIF2023-008738 to J.S.), the Brotman Baty Institute for Precision Medicine and a grant from the National Institutes of Health (R01HG010632 to J.S.). A.V.M. was supported in part by Public Health Service, National Research Service Award, T32GM007270, from the National Institute of General Medical Sciences. C.G.B. was supported in part by a Public Health Service, National Research Service Award, T32HG000035, from the National Human Genome Research Institute. J.-B.L. was supported by a Damon Runyon Cancer Research Foundation fellowship (DRG-2435-21) and a Next-Generation Scientist award from the Cancer Research Society (grant no. 1155581). H.K. is a Washington Research Foundation Postdoctoral Fellow. J.S. is an Investigator of the Howard Hughes Medical Institute.

## AUTHOR CONTRIBUTIONS

A.V.M., B.K.M. and J.S. conceptualized LAMPRA. A.V.M. and B.K.M. designed all assays and carried out all cloning steps. A.V.M. and C.G.B. carried out all cell culture experiments. B.K.M. carried out all library preparation and short-read sequencing. H.K. provided assistance with PacBio sequencing and interpretation. J.-B.L and T.L. wrote computational scripts for calculating MPRA activity scores. C.G.B. and A.V.M. analyzed results with inputs from J.-B.L, T.L., B.K.M. and J.S. A.V.M., C.G.B. and J.S. wrote the initial draft of the manuscript with input from all other authors. A.V.M. and J.S. wrote subsequent drafts of the manuscript with input from all other authors. J.S. supervised the project.

## AI DISCLOSURE STATEMENT

We disclose that data exploration, data analysis, coding and manuscript writing were supported by AI-based tools. The authors take full responsibility for the data, code, analyses, conclusions and writing.

## SUPPLEMENTARY TABLES

**Supplementary Table 1**. NMU Regulatory Element Sequences

**Supplementary Table 2**. Oligonucleotides

## DATA & CODE AVAILABILITY

Code and scripts used for analyses, plasmid maps, and custom sequencing amplicon structures have been deposited on GitHub at https://github.com/shendurelab/LAMPRA. Raw and processed sequencing data generated in this study have been deposited to the Gene Expression Omnibus (GEO, accession number GSE341622).

---

**Supplementary Figure 1.**
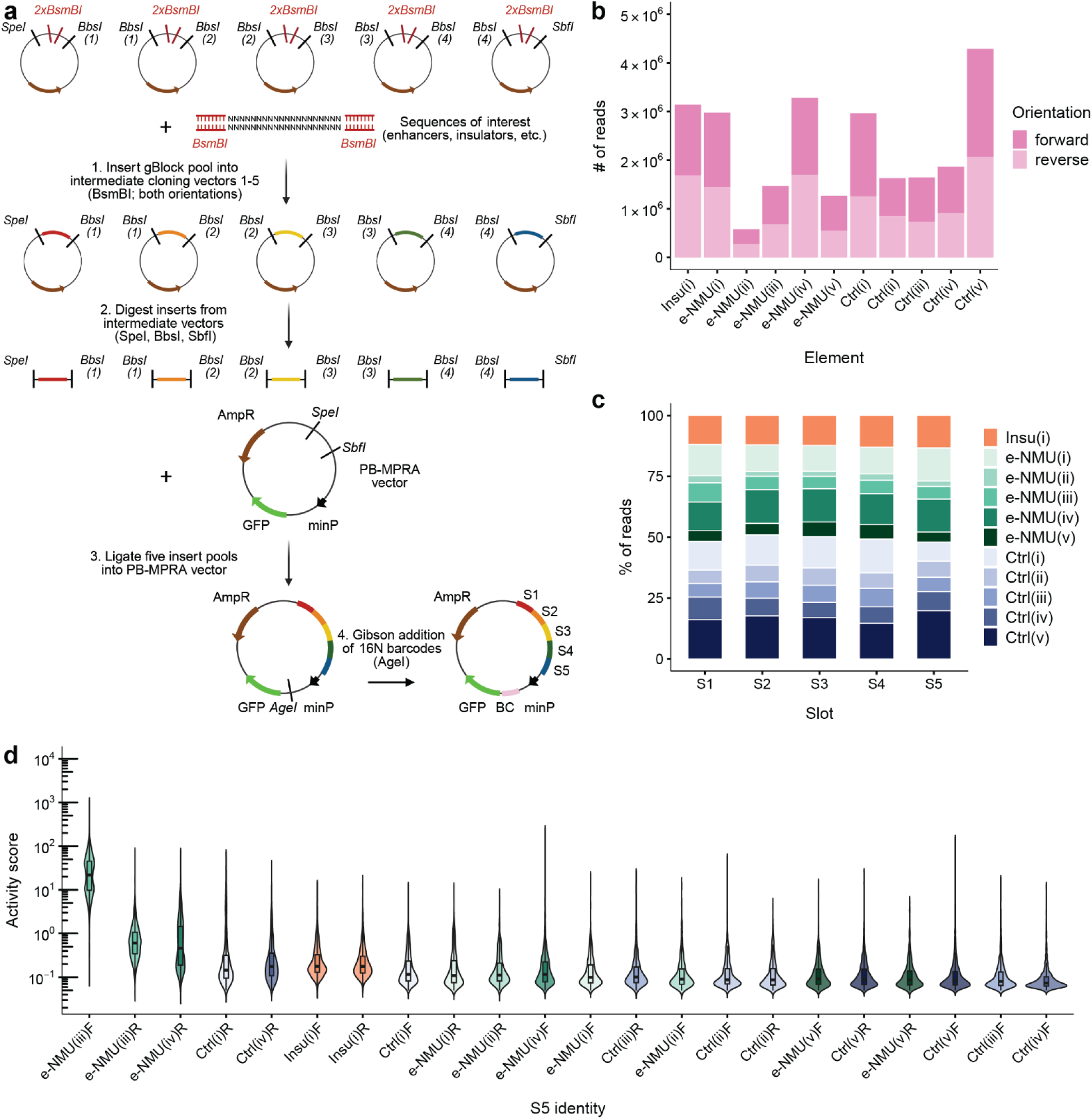
e-NMU LAMPRA library construction, composition, and S5-stratified activity. **(a)** Schematic of key steps of e-NMU LAMPRA library construction. Elements are independently cloned into slot-specific vectors (step 1), combinatorially assembled upstream of a minimal promoter-GFP reporter cassette (steps 2 & 3), and uniquely tagged by degenerate barcodes positioned downstream of the minimal promoter (step 4). **(b)** Stacked bar plot of the number of PacBio reads in which each of the 11 elements appears in any of the 5 slots (S1-S5) of the plasmid library. Each bar is partitioned into two segments by element orientation relative to the minimal promoter. **(c)** Stacked bar plot showing, for each slot (S1-S5) of the plasmid library, the percentage of reads contributed by each element. Each bar sums to 100% and is colored by element identity, collapsing both orientations. **(d)** Violin plots of the distribution of activity scores (y-axis, log10 scale) for sCRLs stratified by S5 identity, sorted by descending mean activity. Overlaid boxes denote the median and interquartile range; whiskers extend to 1.5× IQR.

**Supplementary Figure 2.**
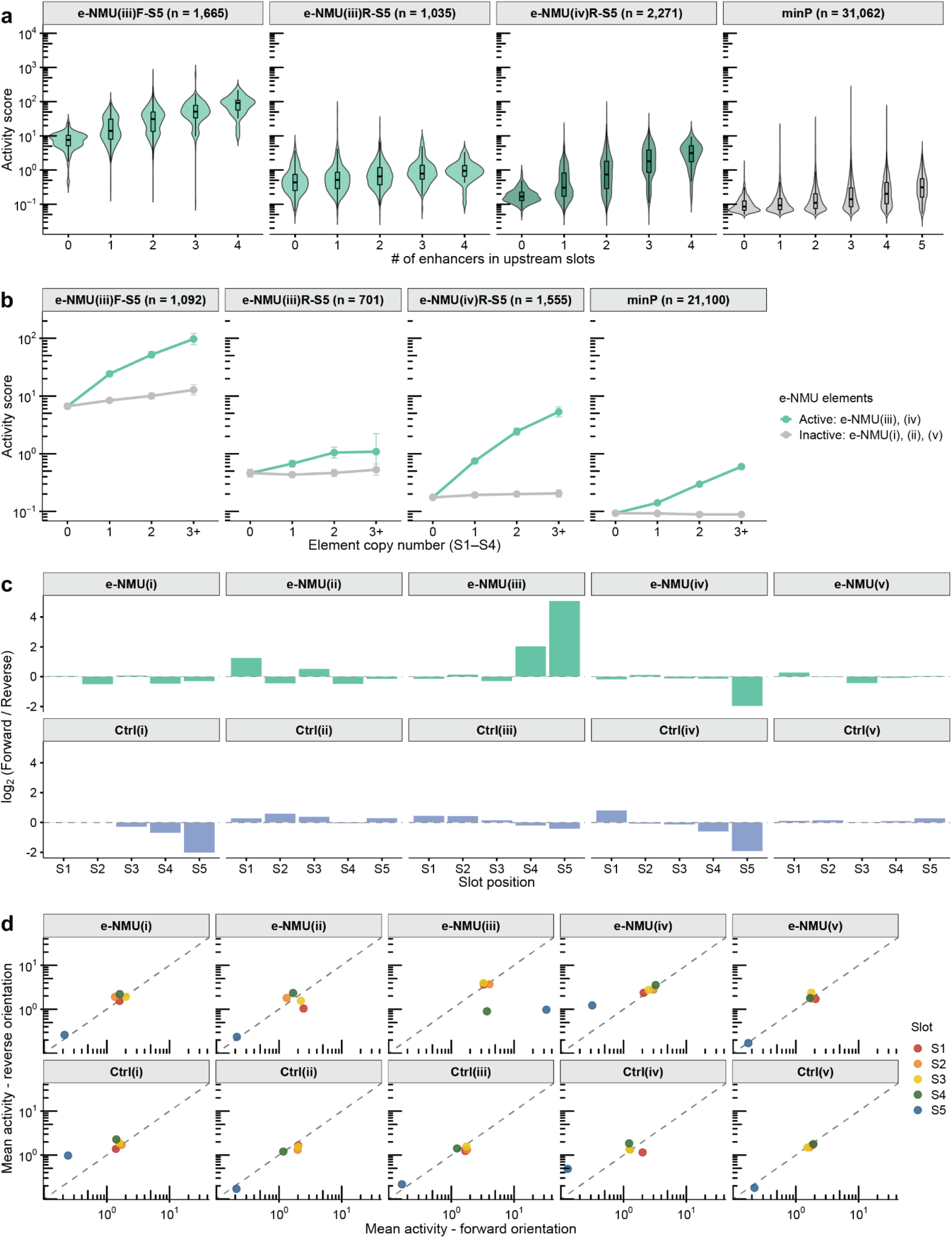
e-NMU enhancer activity scales with copy number and, in S5 only, depends on orientation. **(a)** Violin plots of the distribution of activity scores (y-axis, log10 scale) as a function of the number of e-NMU enhancer elements upstream of the effective promoter (n = 36,033 sCRLs). Elements are stratified to each of four promoter groups. Overlaid boxes denote the median and interquartile range; whiskers extend to 1.5× IQR. **(b)** Mean sCRL activity score as a function of the number of active (e-NMU(iii) and (iv)) or inactive (e-NMU(i), (ii), and (v)) enhancer elements upstream of the promoter, faceted by promoter group. The two line plots are built from non-overlapping sCRLs: the "active" line includes only sCRLs with zero inactive elements in S1-S4, and the "inactive" line includes only sCRLs with zero active elements in S1-S4; sCRLs containing a mix of both are excluded here. Error bars, 95% confidence intervals (t-distribution). **(c)** Bar plots of log₂(forward/reverse) fold-change in mean activity score (y-axis) by slot position S1-S5, faceted by element identity: e-NMU(i)-(v) in the top row, Ctrl(i)-(v) in the bottom row. Dashed line at 0 corresponds to no fold-change. e-NMU(iii) and e-NMU(iv) show orientation-independent activity in upstream slots but a strong orientation bias in S5 (and, for e-NMU(iii), in S4, consistent with promoter activity strong enough to drive detectable transcription even at that distance from the barcode). **(d)** Scatter plots of marginal mean activity in the forward vs. reverse orientation for each element at each slot position (S1-S5), colored by slot and faceted by element identity: e-NMU(i)-(v) in the top row, Ctrl(i)-(v) in the bottom row. Points falling on the dashed identity line (y = x) indicate orientation-independent activity, while distance from the line reflects the degree of orientation dependence. Points are computed by pooling all constructs carrying the indicated element/slot/orientation and averaging activity scores (n = 36,033 sCRLs).

**Supplementary Figure 3.**
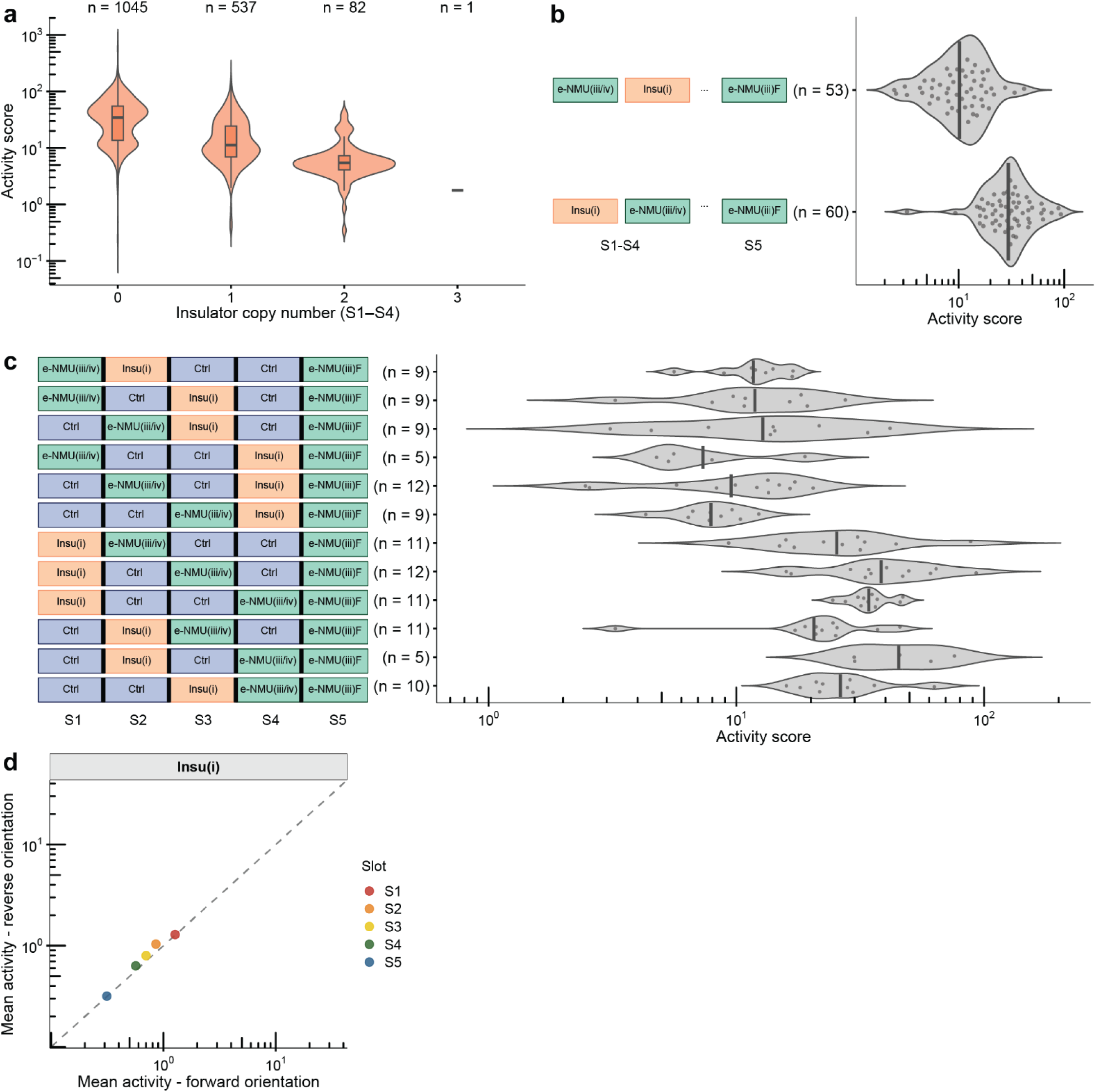
The synthetic insulator suppresses activity when interposed between enhancer and promoter. **(a)** Violin plots of the distribution of activity scores (y-axis, log10 scale) for sCRLs stratified by their copy number of Insu(i) elements across slots S1-S4 (e-NMU(iii)F-S5 subset, n = 1,665 sCRLs). Overlaid boxes denote the median and interquartile range; whiskers extend to 1.5× IQR. **(b)** Violin plots of mean activity score (x-axis, log10 scale) for sCRLs with a single active e-NMU element (e-NMU(iii) or (iv), in either orientation) positioned either upstream or downstream of a single Insu(i) element within S1-S4 (e-NMU(iii)F-S5 subset; sCRLs with any inactive e-NMU element in S1-S4 excluded, n = 113). Points show individual sCRL activity scores, vertical lines denote the mean. **(c)** Similar to panel **b**, but here broken down by full sCRL architecture. Diagrams on the left indicate the position of the Insu(i) and the active e-NMU element within slots S1-S4. **(d)** Scatter plot of marginal mean activity in the forward vs. reverse orientation for Insu(i) at each slot position (S1-S5), colored by slot. Points falling on the dashed identity line (y = x) indicate orientation-independent activity, while distance from the line reflects the degree of orientation dependence. Points are computed by pooling all constructs carrying the indicated element/slot/orientation and averaging activity scores (n = 16,953 sCRLs).

## ONLINE METHODS

### Regulatory element design

Five enhancer regions corresponding to e-NMU regions i-v from Gasperini *et al*.^11^ were selected by centering 1-kb regions on each peak, shifting as necessary to avoid overlapping tiles. Any BsmBI or BbsI restriction sites were removed by mutating one base. To generate corresponding dinucleotide-shuffled negative control sequences for each enhancer region, we used Fasta-uShuffle^51,52^ with the -k 2 option to preserve dinucleotide frequencies while randomizing sequence order. For the synthetic insulator, previously identified insulator elements A2, A4, B2, C3, and F1 from Liu *et al*.^43^ were concatenated with random DNA stuffer sequences to achieve a total length of 1 kb. All 11 designed regions were synthesized as gBlocks (IDT, **Supplementary Table 1**) with flanking BsmBI sites and overhangs (left adaptor: 5’-GATGCGTCCGTCTCACATG-3’ and right adaptor: 5’-CATGTGAGACGACTGTACT-3’) for bidirectional assembly into five intermediate vectors. gBlocks were resuspended to 10 ng/μl and pooled in equal proportions.

### Plasmid library cloning

Five intermediate vectors containing Golden Gate-compatible overhangs were created by digesting the piggyBac-MPRA vector (PB-MPRA) with SpeI-HF (NEB, R3133S) and SbfI-HF (NEB, R3642S), followed by insertion of cloning adaptors. The adaptors were ordered as oligos (IDT, oAVM_081-085, **Supplementary Table 2**) and made double-stranded using oAVM_086 before insertion using NEBuilder HiFi DNA Assembly (NEB, E2621L). Both the intermediate vectors and the pooled gBlocks were digested with BsmBI-v2 (NEB, R0739S) and the vectors were additionally dephosphorylated with Antarctic Phosphatase (NEB, M0289S). All fragments were gel purified and then ligated together in 5 separate ligation reactions. The ligations were then transformed into electrocompetent cells (NEB, C3020K) and plasmid DNA was isolated using the Chargeswitch Midiprep Kit (ThermoFisher, CS31104).

To assemble the final pooled library, the 5 intermediate libraries were digested as follows: intermediate library 1 with SpeI-HF/BbsI, intermediate libraries 2-4 with BbsI, and intermediate library 5 with BbsI/SbfI-HF. The digests were gel purified, yielding gBlock inserts with compatible sticky ends for ordered assembly into the final vector. These five purified inserts were then cloned into the PB-MPRA backbone that had been digested with SpeI-HF and SbfI-HF. Ligation reactions contained 50 ng backbone, 8.8 ng of each of the 5 inserts, 3 µL 10X T4 ligase buffer, 2 µL T4 ligase (NEB, M0202S), and nuclease-free water to a final volume of 30 µL. Before adding the ligase, the reaction was warmed to 37 °C for 2 min and allowed to cool. Ligase was then added and the reaction was incubated at 16 °C overnight. The ligation was transformed into electrocompetent cells (NEB, C3020K), and 50 µL of the 1 mL transformation was plated to count colonies; the remaining transformation was midiprepped to isolate the final pooled library plasmid DNA.

Next, the library was barcoded with a 16-mer degenerate barcode. The library was first digested with AgeI-HF (NEB, R3552S). To generate the double-stranded barcode fragment, 1 μL 100 μM rand-mpra-bc-f, 1 μL 100 μM rand-mpra-bc-r, 8 μL 10 mM Tris pH 8.0, and 10 μL OneTaq Master Mix (NEB, M0482S) were mixed. The oligos were extended through a single PCR cycle with 50 °C annealing and the product was purified using the Monarch PCR & DNA Cleanup Kit (NEB, T1130L). The purified double-stranded barcode was then ligated to the digested library using NEBuilder HiFi DNA Assembly (NEB, E2621L). The assembly product was transformed into electrocompetent cells and plasmid DNA was purified by midiprep as described above.

### PacBio library preparation and sequencing for sCRL-barcode association

For PacBio sequencing, hairpin oligonucleotides were prepared by diluting custom oligo pb_SpeI to 20 µM in 10 mM Tris pH 8.0, 0.1 mM EDTA, and 100 mM NaCl. The oligos were heated to 85 °C and immediately snap-cooled on ice. The plasmid DNA library (5 µg) was digested in 50ul total volume with SpeI-HF (NEB, R3133S) at 37 °C for 2 hours without subsequent heat inactivation. For hairpin ligation, 1 µL of hairpin oligo, 1 µL of 10× CutSmart buffer, 6 µL of 10 mM ATP, and 2 µL of T4 DNA ligase (NEB, M0202S) were added and incubated at room temperature for 20 min. Exonuclease treatment was then performed by adding 1 µL each of Exonuclease III (NEB, M0206S) and Exonuclease VII (NEB, M0379S), followed by incubation at 37 °C for 15 min. DNA was purified using a 1.3× ratio of AMPure PB beads (Pacific Biosciences, 100-265-900), eluted in 30 µL of elution buffer (EB), and quantified with Qubit dsDNA HS (Thermo Fisher). Finally, the resulting library was processed using the PacBio annealing, binding, and cleanup (ABC) workflow according to the manufacturer’s protocol and sequenced on the Vega platform for 24 hours.

### Generation of a sCRL-barcode association dictionary from PacBio sequencing

PacBio consensus reads were processed through a series of custom scripts to associate unique barcodes with sCRLs. First, all reads were oriented to the same direction to ensure consistent downstream processing. Next, elements and barcodes were extracted by identifying fiducials – known fixed sequences that serve as positional markers – flanking the insert regions and extracting the downstream sequences following the fiducials.

Fiducial sequences:

Upstream of insert 1: CTAGTCATG
Upstream of insert 2: CATGAGGACATG
Upstream of insert 3: CATGAGCCCATG
Upstream of insert 4: CATGACATCATG
Upstream of insert 5: CATGCACCCATG
Upstream of 16mer barcode: CTCTTCCGATCT
Downstream of 16mer barcode: CCGGTCGCCACC

Element sequences were then annotated by comparing extracted sequences against the designed enhancer, synthetic insulator, and shuffled negative control sequences. Sequences that exactly matched the first 12 bp of a known element were annotated with the corresponding element name and orientation. Random barcode sequences were extracted using the same method, identifying 16 bp sequences between the fiducial sequences flanking the barcode insertion region. Each read was represented as the combination of elements in slots 1-5 and its corresponding barcode, and counts were collapsed by aggregating reads by unique element-barcode combinations to generate a final sCRL-barcode dictionary.

### Cell culture, transfection, and collection

K562 cells (ATCC, CCL-243) were grown in RPMI 1640 medium (Thermo Fisher, 11875119), supplemented with 10% FBS (Fisher Scientific, Cytiva HyClone fetal bovine serum, SH3039603) and 1× penicillin/streptomycin (Thermo Fisher, 15140122). Cells were kept at 37 °C and 5% CO2, and passaged every 2-3 days.

All cells were transfected in mid-exponential phase. K562 cells were nucleofected using a Lonza 4D-Nucleofector following the manufacturer’s protocol (Lonza, SF cell line kit, program FF-120). For each replicate, 2 × 10⁶ cells were transfected with 2 µg of plasmid DNA (a mixture of cargo plasmid and a plasmid expressing the hyperactive transposase hyPBase, at a mass ratio of 2.5:1). Medium was changed the next day, and cells were passaged as usual thereafter for 10 days post-nucleofection to allow complete dilution of nonintegrated plasmids. After 10 days, cells were harvested for genomic DNA and RNA extraction.

### Bulk MPRA library preparation

Genomic DNA and RNA was extracted using the AllPrep DNA/RNA Mini Kit (Qiagen, 80204) with vortexing to lyse cells.

MPRA amplicon libraries from genomic DNA were generated in two steps of PCR amplification with OneTaq (NEB, M0482S). Approximately 10 µg of genomic DNA input was used per replicate. For the first low-cycle PCR (PCR1), gDNA was mixed with 200 µl 2× OneTaq master mix, 2 µl 100 µM indexed P5 primer (rand-P5-E##), 2 µl 100 µM P7-pLSmp-assUMI-gfp primer, and water to a final volume of 400 µl. This mix was prepared for each replicate and then split over 16 wells for PCR. Cycling parameters were: 2 min at 95 °C, followed by three cycles of 20 s at 95 °C, 30 s at 55 °C, and 2 min at 72 °C, with a final extension of 5 min at 72 °C, followed by a 4 °C hold. Primer P7-pLSmp-assUMI-gfp contains ten random nucleotides (Ns) to serve as a pseudo-UMI (hereafter referred to as UMIs for brevity) to correct for PCR jackpotting. Reactions were pooled for each replicate, cleaned up with AMPure XP beads (Beckman Coulter, A63881) at 1.8×, and eluted in 97 µl of 10 mM Tris pH 8.0. A second round of PCR (PCR2) was performed for each replicate with primers P5 and P7, which amplified the library for sequencing without adding additional UMIs. The entire 97 µl of eluate from PCR1 was used as input and combined with 100 µl OneTaq master mix, 1 µl 100× SYBR Green (Thermo Fisher, S7563), 1 µl 100 µM P5 primer, and 1 µl 100 µM P7 primer. This mix was split over 8 reactions for each replicate. Libraries were amplified and monitored in real-time by qPCR with 2 min at 95 °C, followed by cycles up to the qPCR inflection point consisting of 15 s at 95 °C, 20 s at 60 °C, and 2 min at 72 °C. Libraries were then pooled by replicate and cleaned up with AMPure XP beads (Beckman Coulter, A63881) at 1.8×.

Amplicon libraries from RNA were obtained by first DNase-treating the RNA (up to 100 µl total RNA) by adding 0.1 volume 10× TURBO DNase buffer and 0.5 µl TURBO DNase (Thermo Fisher, AM1907), then incubating at 37 °C for 30 min. An additional 0.5 µl of TURBO DNase was then added followed by a second incubation. DNase was removed by adding 0.1 volume of DNase Inactivation Reagent. For each replicate, 40 µg of DNase-treated RNA was then used for eight reverse transcription reactions. For each reaction, 5 µg RNA was mixed with 1 µl 2 µM P7-pLSmp-assUMI-gfp and 1 µl of 10 mM dNTPs (NEB, N0447L), incubated at 65 °C for 5 min, and placed on ice. 6 µl of reverse transcription master mix was then added (4 µl 5× FS buffer, 1 µl 0.1 M dithiothreitol, and 1 µl SSIV (Thermo Fisher, 18091050)), and the reaction was incubated at 55 °C for 60 min, followed by 80 °C for 10 min. Each reverse transcription reaction was combined and split into 32 total PCR reactions per replicate. For PCR1, reactions contained 160 µl template, 640 µl 2× OneTaq master mix, 8 µl 100 µM indexed P5 primer (rand-P5-E##), 8 µl 100 µM P7 primer, and 464 µl water, and were split over 32 wells with the following cycling parameters: 2 min at 95 °C, followed by three cycles of 20 s at 95 °C, 30 s at 55 °C, and 2 min at 72 °C, with a final extension of 5 min at 72 °C, followed by a 4 °C hold. Reactions were pooled by replicate, cleaned up with AMPure XP beads at 1.8×, and eluted in 291 µl of 10 mM Tris pH 8.0. For PCR2, the entire 291 µl of eluate from PCR1 was used as input and combined with 300 µl OneTaq master mix, 3 µl 100× SYBR Green (Thermo Fisher, S7563), 3 µl 100 µM P5 primer, and 3 µl 100 µM P7 primer. This mix was split over 24 reactions for each replicate. Libraries were amplified and monitored in real-time by qPCR with 2 min at 95 °C, followed by cycles up to the qPCR inflection point consisting of 15 s at 95 °C, 20 s at 60 °C, and 2 min at 72 °C. Libraries were then pooled by replicate.

The three DNA libraries were pooled together, as were the three RNA libraries, and these two final pools were size-selected on 1% agarose gel for the expected band (156 bp). The RNA and DNA pooled libraries were quantified with Qubit dsDNA HS (Thermo Fisher) and diluted to 2 nM. Libraries were loaded at a ratio of 2:1 RNA:DNA and paired-end sequencing was performed on a NextSeq 2000 with the following primers and cycle numbers: read1 (BC forward): 16 cycles with primer TruSeq-R1; index1 (UMI): 10 cycles with primer randMPRAseq-I1; read2 (BC reverse): 16 cycles with primer randMPRAseq-R2; index2 (sample index): 10 cycles with primer TruSeq-I2.

### Bulk MPRA data processing and quantification

Sequencing data were demultiplexed using bcl2fastq. Forward and reverse barcode reads were merged and error-corrected using PEAR (v0.9.11) with parameters set to the barcode length (options -v 16 -m 16 -n 16 -t 16). Using custom Python and R scripts, successfully assembled barcode reads were combined with UMI reads, BC-UMI pairs were counted, and the read and UMI counts per BC were determined. The read and UMI counts for barcodes present in the initial plasmid library (determined a priori; see above) were collected for downstream analysis.

Expression for each barcode from the UMI counts table was computed as follows. First, the total UMIs per replicate corresponding to barcodes in our list were determined for both RNA- and DNA-derived libraries. Barcodes with fewer than 2 DNA UMIs were excluded. Each barcode UMI count was then normalized by the sum of counts in its respective sample type (DNA or RNA) and winsorized at the 99th percentile independently for RNA and DNA prior to aggregation. The normalized RNA UMI count was then divided by the normalized DNA UMI count to generate the bulk MPRA-derived estimate of expression per barcode ("activity score"). We then used the a priori sCRL-barcode association to assign activity scores to each sCRL, filtered for sCRLs present in 2 or more replicates, and averaged activity scores across replicates to obtain a single activity score per sCRL. For the replicate correlation analysis and all activity score visualizations, a pseudocount of 0.05 was added to the scores prior to log-transformation to retain sCRLs with no detectable activity (*i.e.* zero values).

### Regression to predict LAMPRA activity scores

Replicate-averaged LAMPRA activity scores were modeled as a function of sCRL composition. Each slot in the five-slot assembly was described by two categorical variables (5 x 2 = 10 features): (a) element identity (11 levels) and (b) element orientation (2 levels). For linear models, which cannot accommodate integer-encoded categories, these features were one-hot encoded, with the first variable dropped to avoid perfect multicollinearity (n = 55 features without interaction terms; n = 1,485 features with interaction terms). For ensemble methods, the categorical representations were retained (n = 10 features).

Of sCRLs present in two or more replicates (n = 36,033), 15% were randomly sampled for model evaluation (n = 5,405). The remaining observations were randomly split 90:10 into training and validation sets of 27,565 and 3,063, respectively. Ordinary least squares (OLS), Lasso, Ridge, and ElasticNet linear models were fit and, where applicable, hyperparameters were tuned to minimize mean squared error on the validation set. The best-performing models by this metric were evaluated on the held-out test set. Random forest (RF) and gradient-boosted regressors were constructed and evaluated in the same manner. The linear models and the random forest model were implemented with scikit-learn (v1.6.1), while gradient-boosted trees were implemented with XGBoost (v3.1.2). The exact hyperparameters searched in regularized linear model and ensemble model training can be found at: https://github.com/shendurelab/LAMPRA.

### Downsampling analysis

The RF model was re-instantiated with fixed hyperparameters and re-fit on randomly downsampled subsets of the training data. To preserve exact correspondence with the original split, only the original training set (27,565 sCRLs) was subsampled, and the validation set (3,063 sCRLs) was excluded. Within a subsampling replicate, the training set was varied across 100 nested fractions, from 1% to 100% of the training data in 1% increments (276 to 27,565 sCRLs). Model performance was evaluated on the same held-out test set (5,405 sCRLs) for each fraction. This procedure was repeated and performance averaged across 50 subsampling replicates.

